# Linkage disequilibrium connects genetic records of relatives typed with disjoint genomic marker sets

**DOI:** 10.1101/345322

**Authors:** Jaehee Kim, Michael D. Edge, Bridget F. B. Algee-Hewitt, Jun Z. Li, Noah A. Rosenberg

## Abstract

In familial searching in forensic genetics, a query DNA profile is tested against a database to determine whether it represents a relative of a database entrant. We examine the potential for using linkage disequilibrium to identify pairs of profiles as belonging to relatives when the query and database rely on nonoverlapping genetic markers. Considering data on individuals genotyped with both microsatellites used in forensic applications and genome-wide SNPs, we find that ~30-32% of parent–offspring pairs and ~35-36% of sib pairs can be identified from the SNPs of one member of the pair and the microsatellites of the other. The method suggests the possibility of performing familial searches of microsatellite databases using query SNP profiles, or vice versa. It also reveals that privacy concerns arising from computations across multiple databases that share no genetic markers in common entail risks not only for database entrants, but for their close relatives as well.

## Introduction

Forensic DNA testing sometimes seeks to identify unknown individuals through familial searching, or relatedness profiling. When no exact match of a query DNA profile to a database of profiles is found, investigators can potentially test for a partial match to determine whether the query profile might instead represent a close relative of a person whose profile appears in the database [1–3]. A positive test leads investigators to consider relatives of the person with the partial match as possible contributors of the query profile.

Familial searching expands the potential to identify unknown contributors beyond the level achieved when searching exclusively for exact database matches. The larger set of people accessible to investigators—database entrants, plus their relatives—can increase the probability that the true contributor of a query profile is identified [1, 4]. However, the accessibility of relatives to investigators in database queries raises privacy and legal policy concerns, as considerations guiding appropriate inclusion of DNA profiles in databases and subsequent use of those profiles generally focus on the contributors of the profiles rather than on the close relatives potentially accessible to investigators from those profiles [5, 6]. Concerns about privacy vary in magnitude across populations, as false positive identifications of relatives might be substantially more likely to affect members of populations with lower genetic diversity and hence a greater likelihood of chance partial matches [7, 8], or members of populations overrepresented in DNA databases [5, 9].

Recently, considering genome-wide single-nucleotide polymorphisms (SNPs) together with the Combined DNA Index System (CODIS) microsatellite markers used for forensic genetic databases in the United States elsewhere [10–12], we studied the possibility of matching a forensic-genetic record in one database to a profile on a set of nonoverlapping genetic markers in a different database. We showed that records could be matched between databases with no overlapping genetic markers, provided that sufficiently strong linkage disequilibrium (LD) exists between markers represented in the two databases [13]. The approach could facilitate development of new SNP-based forensic marker systems that are backward-compatible with the CODIS microsatellites, as it could enable a query SNP profile to be tested for a match to a microsatellite database, or vice versa. It also uncovers privacy concerns, as an individual present in a SNP database—collected for biomedical research, genealogical research, or personal genomics, for example—might be possible to link to a CODIS profile, and vice versa, in a manner not intended in the context of either database examined in isolation. First, a SNP database entrant could become accessible to forensic investigation. Second, although in the United States, the use of forensic genetic markers given protections against unreasonable searches is based partly on a premise that these markers provide only the capacity for identification and do not expose phenotypic information [14–16], phenotypes that are possible to predict from a SNP profile could potentially be predicted from a CODIS profile by connecting the CODIS profile to a SNP profile and then predicting phenotypes from the SNPs.

Does cross-database record matching extend to relatives? In other words, is it possible to identify a genotype record with one set of genetic markers as originating from a relative of the contributor of a genotype record obtained with a distinct, nonoverlapping set of markers? If so, then new marker systems in the forensic context could permit relatedness profiling in a manner that is compatible with existing marker systems, as a profile from a new SNP or DNA sequence system could be tested for relationship matches to existing microsatellite profiles. However, a substantial privacy concern would also be raised, as inclusion in a biomedical, genealogical, or personal genomics dataset could expose relatives of the participant to forensic investigation; moreover, phenotypes of a relative could potentially be identifiable from a forensic profile.

Here, extending the likelihood framework of Edge et al. [13] to accommodate familial relationships, we devise and evaluate an algorithm for using linkage disequilibrium to perform cross-database matching of relatives. We assess the performance of record matching between SNP profiles and microsatellite profiles gathered with distinct marker sets, in the case in which the SNP profile and the microsatellite profile represent distinct but closely related individuals. The results contribute to the evaluation of the genetic privacy of existing forensic marker systems, as well as to assessment of the potential of familial searching with new SNP or DNA sequence marker systems that might be devised in the future.

## Results

### Likelihood method

We consider a dataset that contains *L* microsatellites, or short-tandem-repeat (STR) loci, each surrounded by an associated set of neighboring SNPs. For two individuals *A* and *B*, we denote by *R_Aℓ_* the diploid genotype of individual *A* at STR locus *ℓ* and by *S_Bℓ_* the set of diploid genotypes of individual *B* at the neighboring SNP loci associated with STR locus *ℓ*. Considering all *L* STR loci, we let *R_A_* be the STR profile of individual *A* from the STR dataset, *R_A_* = {*R_A_*_1_*, R_A_*_2_*,…,R_AL_*}, and we let *S_B_* be the SNP profile of individual *B* from the SNP dataset, *S_B_* = {*S_B_*_1_*, S_B_*_2_*,…, S_BL_*}.

Any familial relationship between individuals *A* and *B* can be completely characterized by nine condensed identity coefficients, Δ_1_, Δ_2_*,…,* Δ_9_, corresponding to the probabilities of the nine possible condensed identity states *C*_1_*, C*_2_*,…, C*_9_ [17, 18] (Table 1). Each *C_i_* represents a possible identity-by-descent configuration of the four alleles in the unordered diploid genotypes of two individuals at an autosomal locus.

**Table 1:**
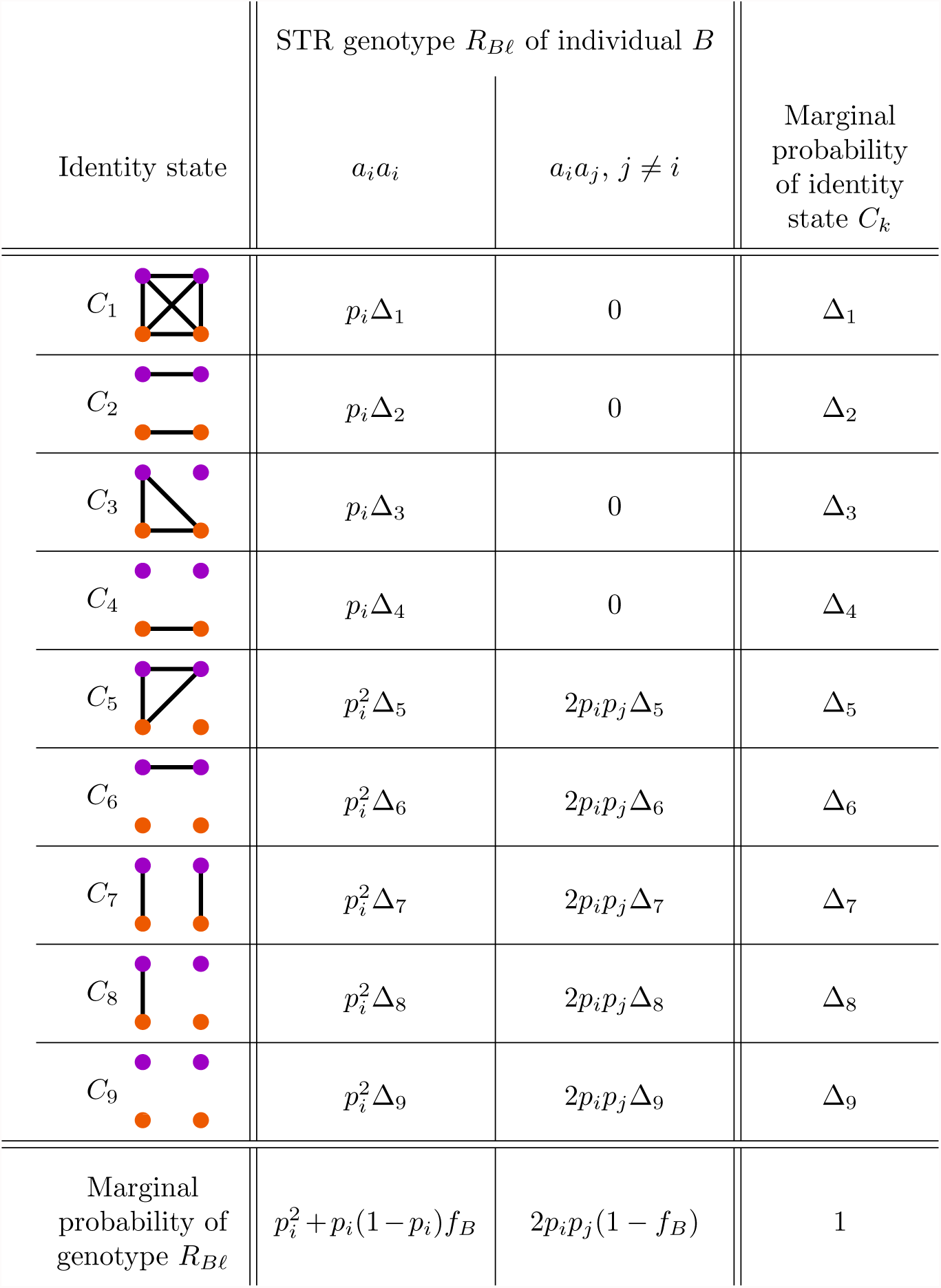
The joint distribution of the condensed identity state *C_k_* and the genotype *R_Bℓ_* of individual *B* at STR locus *ℓ*. In the nine condensed identity states *C*_1_ to *C*_9_ for individuals *A* and *B*, the alleles of individuals *A* and *B* appear in purple and orange, respectively. Identical-by-descent alleles are connected by black lines. The probability of state *C_k_* is denoted Δ*_k_*, with 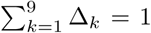. Genotypes are unordered, and *p_i_* denotes the frequency of allele *a_i_*. The quantity *f_B_* represents the inbreeding coefficient of individual *B*, *f_B_* = Δ_1_ + Δ_2_ + Δ_3_ + Δ_4_. The marginal probability of state *C_k_*, 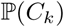, is obtained by summing over all possible alleles *a_i_* and *a_j_* at the locus, 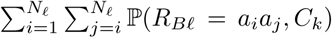, where *N_ℓ_* is the number of distinct alleles at STR locus *ℓ*. The marginal probability of genotype *R_Bℓ_*, 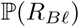, is a sum over identity states: 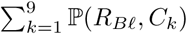.

| Identity state | STR genotype $R_{B\ell}$ of individual $B$ | | Marginal probability of identity state $C_k$ |
| --- | --- | --- | --- |
| | $a_i a_i$ | $a_i a_j, j \neq i$ | |
| $C_1$ | $p_i \Delta_1$ | 0 | $\Delta_1$ |
| $C_2$ | $p_i \Delta_2$ | 0 | $\Delta_2$ |
| $C_3$ | $p_i \Delta_3$ | 0 | $\Delta_3$ |
| $C_4$ | $p_i \Delta_4$ | 0 | $\Delta_4$ |
| $C_5$ | $p_i^2 \Delta_5$ | $2p_i p_j \Delta_5$ | $\Delta_5$ |
| $C_6$ | $p_i^2 \Delta_6$ | $2p_i p_j \Delta_6$ | $\Delta_6$ |
| $C_7$ | $p_i^2 \Delta_7$ | $2p_i p_j \Delta_7$ | $\Delta_7$ |
| $C_8$ | $p_i^2 \Delta_8$ | $2p_i p_j \Delta_8$ | $\Delta_8$ |
| $C_9$ | $p_i^2 \Delta_9$ | $2p_i p_j \Delta_9$ | $\Delta_9$ |
| Marginal probability of genotype $R_{B\ell}$ | $p_i^2 + p_i(1 - p_i)f_B$ | $2p_i p_j(1 - f_B)$ | 1 |

We test a hypothesis that *A* and *B* are related with a relationship defined by a condensed identity coefficient vector, **Δ** = (Δ_1_, Δ_2_*,…,*Δ_9_), against the null hypothesis in which the two individuals are unrelated. To test the hypothesis, we generalize the log-likelihood-ratio match score of Edge et al. [13]:

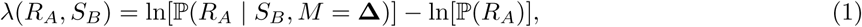
>
where *M* is a variable indicating the hypothesized relationship between individuals *A* and *B*. In the work of Edge et al. [13], *M* was assumed to represent the hypothesis in which *A* and *B* are the same individual. Here, we consider relationship hypotheses more generally. As the natural logarithm of a likelihood ratio, a match score of 10, for example, represents a value of *e*^10^ ≈ 22, 026 for the ratio 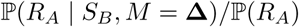.

Assuming independence of the STR loci, so that genotypes at separate STRs are independent, we can rewrite Eq. 1 as a sum of log-likelihood ratios across STR loci:

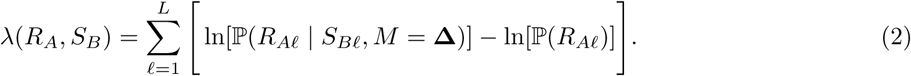

The likelihood 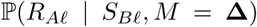 for arbitrary hypotheses **Δ** for the relationship between *A* and *B* is obtained by a decomposition over possible values of *R_Bℓ_*, the STR genotype of *B*. The decomposition, which provides the methodological advance beyond Edge et al. [13], considers products of terms 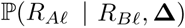 reflecting the relationship of *A* and *B*, and terms 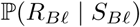, reflecting STR genotype probabilities conditional on surrounding SNP probabilities. Details appear in the Methods (see “Match score calculation”).

### Experimental design

To perform cross-database matching of relatives, we begin from datasets with *N_R_* STR and *N_S_* SNP profiles, where some or all of the profiles in one dataset represent relatives of individuals whose profiles appear in the other dataset. For each pair of profiles (*R_A_, S_B_*), one from each dataset, we compute the match score *λ*(*R_A_, S_B_*) under a specified hypothesis for the relationship **Δ** between the pair. The match-score matrix **X** is an *N_S_* × *N_R_* matrix whose (*i, j*) entry is *λ*(*i, j*). From the match-score matrix, we identify matches according to each of four match-assignment algorithms. For simplicity, we assume *N_R_* = *N_S_*.

### Relationship schemes

We used datasets containing genotypes at 13 STR loci and 642,563 SNP loci in 872 Human Genome Diversity Panel individuals (Methods, “Data”). Although our approach applies for arbitrary relationship hypotheses, we focused on close relationships, for which SNPs of one individual in a relative pair are most likely to contain information about STRs of the other and vice versa. Assuming that individuals were not inbred, we considered three schemes for **Δ**_true_, the true relationship between individuals in the STR and SNP datasets: (1) same individual, **Δ**_true_ = (0, 0, 0, 0, 0, 0, 1, 0, 0); (2) parent–offspring, **Δ**_true_ = (0, 0, 0, 0, 0, 0, 0, 1, 0); and (3) sibling pairs, 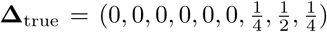 For schemes (2) and (3), we simulated pedigrees based on the actual genotype data to produce datasets containing relatives (Methods, “Pedigree generation”).

Following Edge et al. [13], for each scheme for the true relationship between individuals in the STR and SNP datasets, we generated 100 random partitions of the individuals into a training set (75%) and a test set (25%) (Methods, “Training and test sets”). For each partition, using BEAGLE [19, 20], we phased the training set to produce haplotypes containing STR loci and their surrounding SNPs (Methods, “BEAGLE settings”). Next, as in Edge et al. [13], we used the phased training set as a reference and augmented it with the SNP genotypes of the unphased test set. We then used BEAGLE to impute genotypes at the STR loci in the test set based on the neighboring SNPs (Methods, “Training and test sets”).

Once STR genotype probabilities for individuals in the SNP dataset were obtained by imputation, we computed the match score (Eq. 2) for all possible pairs of individuals, one in the SNP dataset and one in the STR dataset, under three relationship hypotheses: (1) same individual, (2) parent–offspring, and (3) sib pairs. Letting **Δ**_test_ denote the hypothesized relationship from which we computed the match score, we considered nine choices for the pair (**Δ**_true_, **Δ**_test_). A schematic of the computation appears in Figure 1.

**Figure 1:**
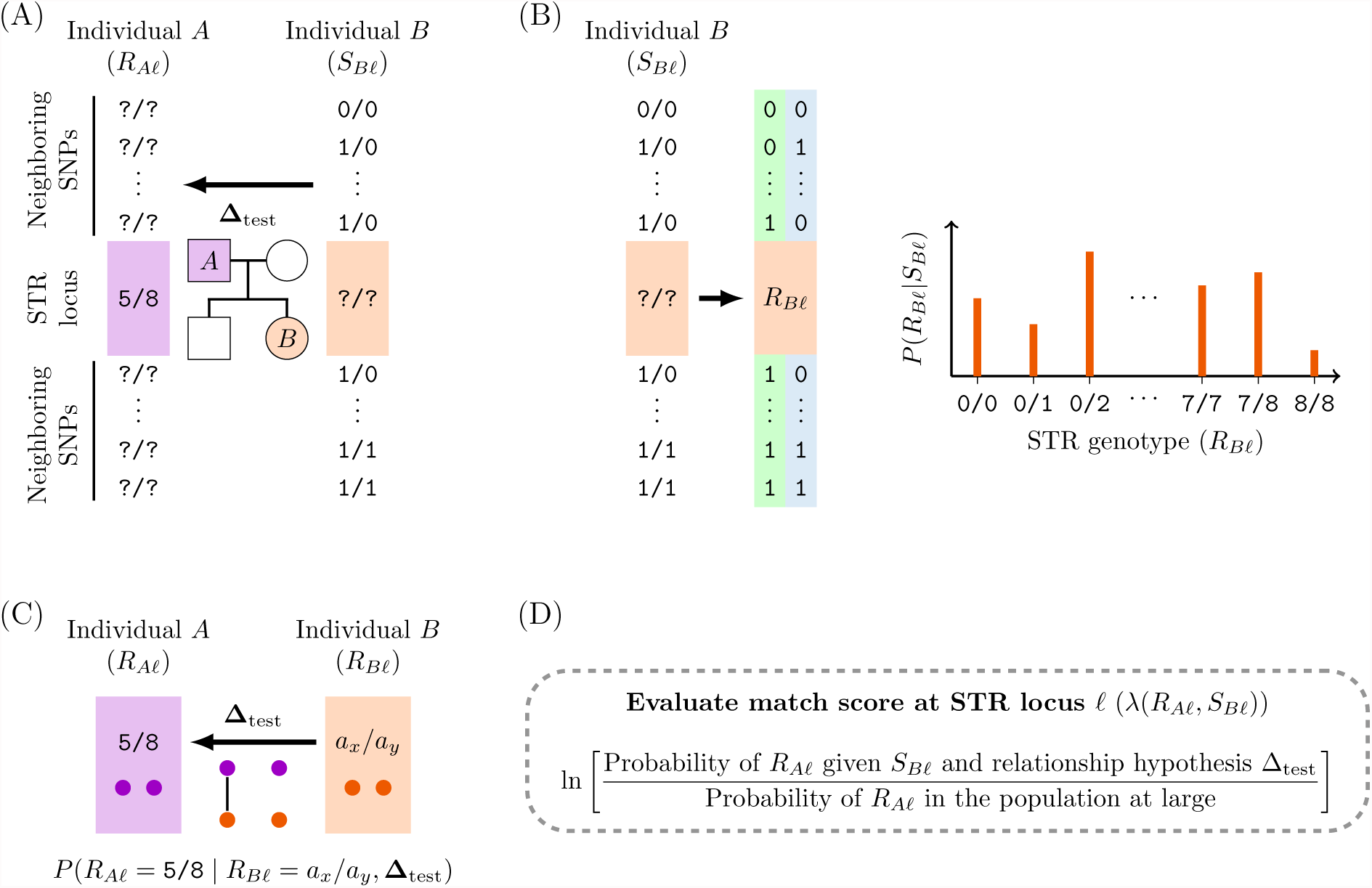
Schematic for evaluating *λ*(*R_A_, S_B_*), the match score for a pairing of STR profile *R_A_* of individual *A* and SNP profile *S_B_* of individual *B*, assuming a relationship hypothesis **Δ**_test_ between individuals *A* and *B*. Because the match score is a sum of contributions from *L* STR loci (Eq. 2), we illustrate steps for computing the match score *λ*(*R_Aℓ_, S_Bℓ_*) of a single locus *ℓ*. (A) The data and the hypothesis tested. Consider individual *A* with unordered diploid STR genotype *R_Aℓ_* = 5/8 and individual *B* with SNP profile *S_Bℓ_* around locus *ℓ*. We seek to test a hypothesis that *A* is a parent of *B*. (B) Imputation of the STR in individual *B*. Given the SNP profile *S_Bℓ_* of individual *B*, we use `BEAGLE` to estimate the STR genotype probabilities 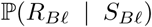 using a phased training set as a reference panel. (C) Conditional probability of the STR genotype of *A* given the probabilistically imputed STR genotype of *B* and a test hypothesis **Δ**_test_. Under the hypothesis **Δ**_test_, considering all condensed identity states possible for a pair of individuals given **Δ**_test_, we compute the probability that individual *A* has the known STR genotype 5/8 conditional on the imputed STR genotype *a_x_/a_y_*: 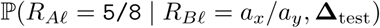. We evaluate this probability for all STR genotypes possible for individual *B*. (D) The match score at locus *ℓ*. Multiplying terms 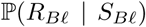 from (B) and 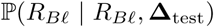 from (C) and summing over all possible STR genotypes *R_Bℓ_* of *B* at locus *ℓ* (Eq. 5), we obtain 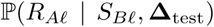, the probability of the STR genotype of individual *A* at locus *ℓ* given the SNP genotype of individual *B* around locus *ℓ* and the relationship hypothesis. The match score of locus *ℓ* (the summand in Eq. 2) is expressed as a log-likelihood ratio of the test hypothesis **Δ**_test_ and the null hypothesis that *A* and *B* are unrelated.

When the relationship hypothesis tested and the true relationship are the same (**Δ**_true_ = **Δ**_test_), diagonal entries in the match-score matrix **X** represent match scores for true relationship matches, and off-diagonal entries give match scores for unrelated individuals. When **Δ**_true_ ≠ **Δ**_test_, however, the model is misspecified; in this case, a diagonal entry gives the match score for the test hypothesis when two individuals are related but the relationship hypothesis tested differs from the true relationship. An off-diagonal entry gives the match score for the test hypothesis for unrelated individuals.

### Match-assignment scenarios

Given a match-score matrix **X** for the pair (**Δ**_true_, **Δ**_test_), we considered four matching scenarios: one-to-one matching, one-to-many matching with a query SNP profile, one-to-many matching with a query STR profile, and needle-in-haystack matching [13]. In one-to-one matching, we assume that it is already known that each profile in the SNP dataset has exactly one true relative in the STR dataset and vice versa. To find the pairing of profiles that maximizes the sum of match scores across all paired profiles, we use the Hungarian algorithm [21] as in Edge et al. [13]. Record-matching accuracy is the fraction of pairs correctly matched.

In one-to-many matching, we relax the one-to-one correspondence and examine the possibility that an observation in one dataset might be identified as having multiple relationship matches in another dataset. In the case in which a SNP profile is used as a “query,” for a given SNP profile, the STR profile with the largest match score is proposed as its match. In other words, for each row of the match-score matrix, we select the largest entry in the row. Record-matching accuracy is the fraction of SNP profiles matched to the correct STR profiles. In this scenario, a SNP profile has exactly one putative STR profile, but an STR profile can be chosen as the match for multiple SNP profiles. Similarly, in one-to-many matching with an STR profile as the query, we select the SNP profile with the largest match score for the given STR profile: for each column of the match-score matrix, the largest entry is chosen as a match. Record-matching accuracy is the fraction of STR profiles matched to the correct SNP profiles.

In needle-in-haystack matching, unlike in the other scenarios, we investigate a setting in which a database query is performed for only one profile. A perfect matching is achieved when no overlap occurs in the match-score distributions of the correct relationship matches and the incorrect matches. We quantified the record-matching accuracy as the proportion of true relatedness matches with greater match scores than the largest match score across all non-matching pairs.

### Record-matching accuracy

For each of the three choices of **Δ**_true_, we considered 100 partitions of the initial sample into a training set and a test set (Methods, “Training and test sets”). For each of the three test hypotheses **Δ**_test_, we then computed the match score (Eq. 2) for all pairs of profiles in the test set, one from the STR dataset and one from the SNP dataset. For each pair (**Δ**_true_, **Δ**_test_), the match-score matrix corresponding to the median record-matching accuracy across 100 partitions in one-to-one matching appears in Figure 2. Table 2 presents the median, minimum, and maximum accuracies.

**Figure 2:**
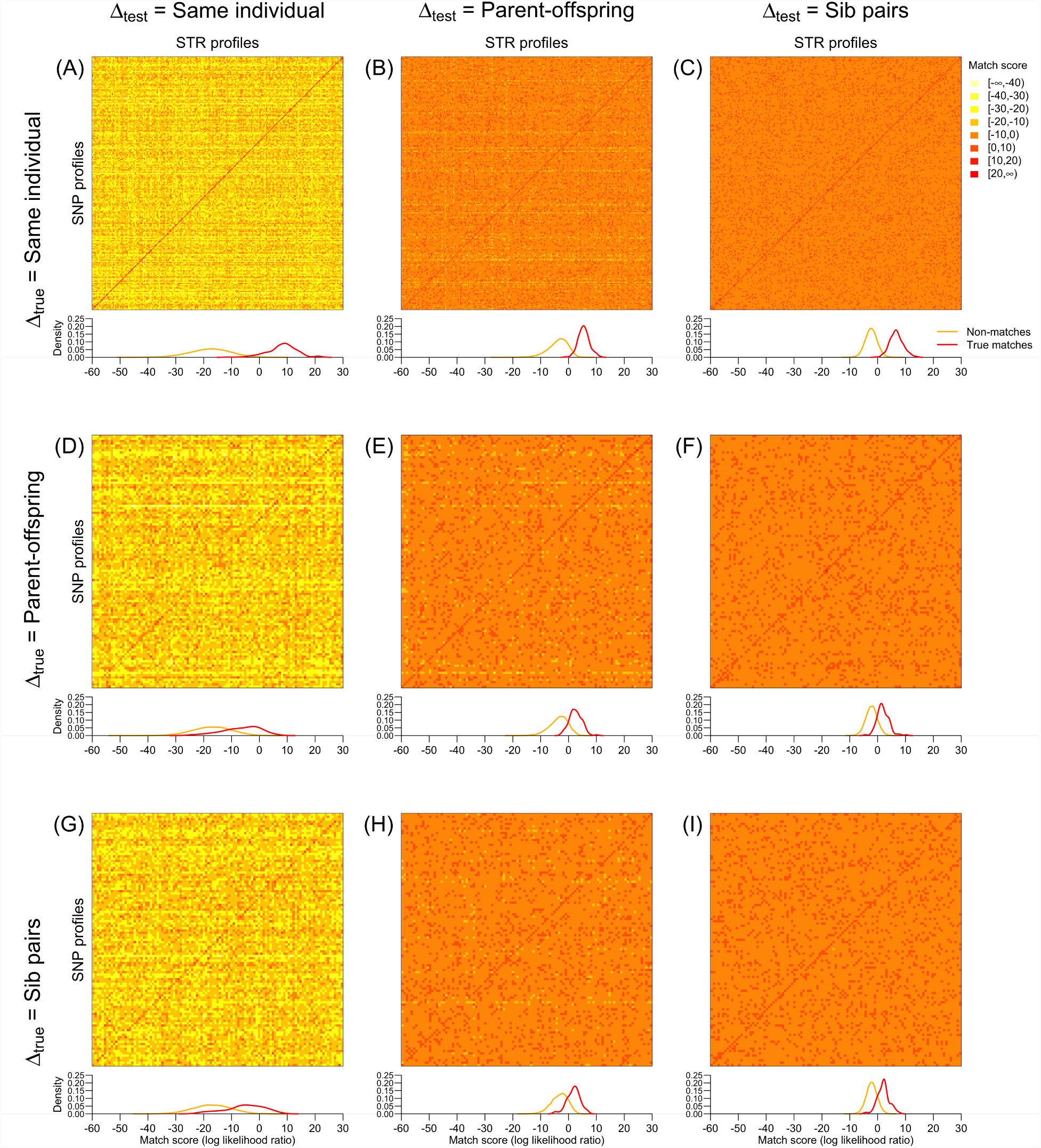
Match scores of SNP and STR profiles in various relatedness settings. **Δ**_true_ denotes a true relationship between pairs of individuals represented on the diagonal of a matrix, one from a SNP dataset and the other from an STR dataset. **Δ**_test_ represents a test relationship hypothesis from which match scores are computed. Of 100 random partitions of individuals into training and test sets, the results from the partition with median one-to-one record-matching accuracy are shown. For each (**Δ**_true_, **Δ**_test_) scheme, we plot the match-score matrix (top) and the kernel density estimate for match scores (bottom). The kernel density estimates separately consider the diagonal entries (true matches) and the off-diagonal entries (non-matches). (A) **Δ**_true_, same individual; **Δ**_test_, same individual. (B) **Δ**_true_, same individual; **Δ**_test_, parent–offspring. (C) **Δ**_true_, same individual; **Δ**_test_, sib pairs. (D) **Δ**_true_, parent–offspring; **Δ**_test_, same individual. (E) **Δ**_true_, parent–offspring; **Δ**_test_, parent–offspring. (F) **Δ**_true_, parent–offspring; **Δ**_test_, sib pairs. (G) **Δ**_true_, sib pairs; **Δ**_test_, same individual. (H) **Δ**_true_, sib pairs; **Δ**_test_, parent–offspring. (I) **Δ**_true_, sib pairs; **Δ**_test_, sib pairs.

**Table 2:** Record-matching accuracies between genome-wide SNP and CODIS STR profiles. For each of three choices for the true relationship between corresponding profiles in the SNP and STR datasets (**Δ**_true_) and for each of three choices for the relationship hypothesis tested (**Δ**_test_), in each of four match-assignment scenarios (one-to-one, one-to-many with a query SNP profile, one-to-many with a query STR profile, and needle-in-haystack), the minimum, median, and maximum accuracies across 100 partitions of the sample into a training set (75%) and test set (25%) are shown. For **Δ**_true_ = **Δ**_test_ (block-diagonal entries), each entry shows the fraction of individuals correctly matched to their true relatives. For **Δ**_true_ ≠ **Δ**_test_, each entry represents the fraction of individuals matched to their true relatives under a misspecified relationship hypothesis **Δ**_test_.

| $\Delta_{\text{true}} \backslash \Delta_{\text{test}}$ | Same individual | | Parent-offspring | | Sib pairs | | Match-assignment scenario |
| --- | --- | --- | --- | --- | --- | --- | --- |
|  | Median | Min, Max | Median | Min, Max | Median | Min, Max |  |
| Same individual | 0.982 | 0.917, 1.000 | 0.885 | 0.807, 0.982 | 0.968 | 0.908, 1.000 | One-to-one |
|  | 0.908 | 0.862, 0.950 | 0.780 | 0.716, 0.858 | 0.878 | 0.830, 0.927 | One-to-many: SNP query |
|  | 0.904 | 0.830, 0.945 | 0.780 | 0.716, 0.844 | 0.688 | 0.596, 0.748 | One-to-many: STR query |
|  | 0.431 | 0.064, 0.697 | 0.085 | 0.000, 0.294 | 0.202 | 0.018, 0.440 | Needle-in-haystack |
| Parent-offspring | 0.174 | 0.101, 0.257 | 0.312 | 0.239, 0.440 | 0.266 | 0.165, 0.367 | One-to-one |
|  | 0.183 | 0.083, 0.266 | 0.303 | 0.229, 0.422 | 0.266 | 0.183, 0.330 | One-to-many: SNP query |
|  | 0.165 | 0.064, 0.239 | 0.321 | 0.220, 0.385 | 0.275 | 0.174, 0.358 | One-to-many: STR query |
|  | 0.000 | 0.000, 0.055 | 0.018 | 0.000, 0.092 | 0.018 | 0.000, 0.092 | Needle-in-haystack |
| Sib pairs | 0.294 | 0.165, 0.450 | 0.303 | 0.174, 0.431 | 0.349 | 0.229, 0.459 | One-to-one |
|  | 0.284 | 0.220, 0.367 | 0.303 | 0.211, 0.394 | 0.349 | 0.248, 0.459 | One-to-many: SNP query |
|  | 0.275 | 0.183, 0.358 | 0.303 | 0.211, 0.413 | 0.358 | 0.257, 0.450 | One-to-many: STR query |
|  | 0.028 | 0.000, 0.110 | 0.028 | 0.000, 0.119 | 0.046 | 0.000, 0.119 | Needle-in-haystack |

When **Δ**_test_ = (0, 0, 0, 0, 0, 0, 1, 0, 0), our generalized match score in Eq. 2 is equivalent to that of Edge et al. [13] for identifying the same individual in two disjoint datasets. Our results under the setting **Δ**_true_ = **Δ**_test_ = (0,0,0,0,0,0,1,0,0) closely follow Edge et al. [13]. Most match scores for true matches exceed most match scores for non-matches, so that the diagonal entries of the match-score matrix have generally larger values than off-diagonal entries (Figure 2A). In one-to-one matching, among 100 partitions into the training and test sets, the median record-matching accuracy is 214 of 218 (98.2%).

With **Δ**_true_ = **Δ**_test_ = (0, 0, 0, 0, 0, 0, 0, 1, 0), we search for parent–offspring relationships between a SNP profile and an STR profile. Match scores for true relationship matches also generally exceed those for non-matches, so that the distribution of diagonal entries is shifted toward higher values compared to the distribution of off-diagonal entries (Figure 2E). The distinction between diagonal and off-diagonal entries is not as great as when profiles represent the same individual rather than parent–offspring pairs. The median record-matching accuracy for one-to-one matching is 31.2% (34 of 109 individuals).

The case of sib-pair relationships between SNP and STR profiles, 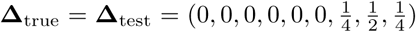 (Figure 2I), is similar to the case of parent–offspring relationships. Diagonal and off-diagonal match-score matrix entries are differentiated, though not as strongly as in matching profiles from the same individual. The median record-matching accuracy for one-to-one matching is 34.9% (38 of 109 individuals).

### Model misspecification

Considering the six (**Δ**_true_, **Δ**_test_) pairs with **Δ**_true_ ≠ **Δ**_test_, we observe that record-matching accuracies for misspecified test hypotheses are generally lower than in corresponding cases with the test hypothesis correctly specified (Figure 2 and Table 2). The value of **Δ**_true_ has a stronger influence on record-matching than does **Δ**_test_; for example, higher accuracies are seen when SNP and STR profiles truly represent the same individual and the test hypothesis is misspecified to search for relative pairs, compared with lower accuracies when profiles represent relatives and the test hypothesis is misspecified to search for exact matches.

For **Δ**_true_ = (0, 0, 0, 0, 0, 0, 1, 0, 0), compared to 98.2% median accuracy when **Δ**_true_ = **Δ**_test_, median accuracy is 88.5% when parent–offspring matches are sought instead of exact matches and 96.8% when sib-pair matches are sought. For **Δ**_true_ = (0, 0, 0, 0, 0, 0, 0, 1, 0), compared to 31.2% median accuracy when **Δ**_true_ = **Δ**_test_, median accuracy is 17.4% for the exact-match test hypothesis and 26.6% for the sib-pair test hypothesis. For 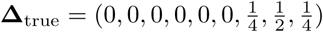, compared to 34.9% median accuracy when **Δ**_true_ =**Δ**_test_, median accuracy is 29.4% when seeking exact matches and 30.3% when seeking parent–offspring matches.

### Match-assignment threshold

The number of false positive matches can be decreased by setting a minimum match-score threshold below which records are left unpaired. For each of the four match-assignment scenarios, the proportions of correct, incorrect, and unassigned profiles with a varying threshold appear in Figure 3 for the partitions with the minimum, median, and maximum record-matching accuracies.

**Figure 3:**
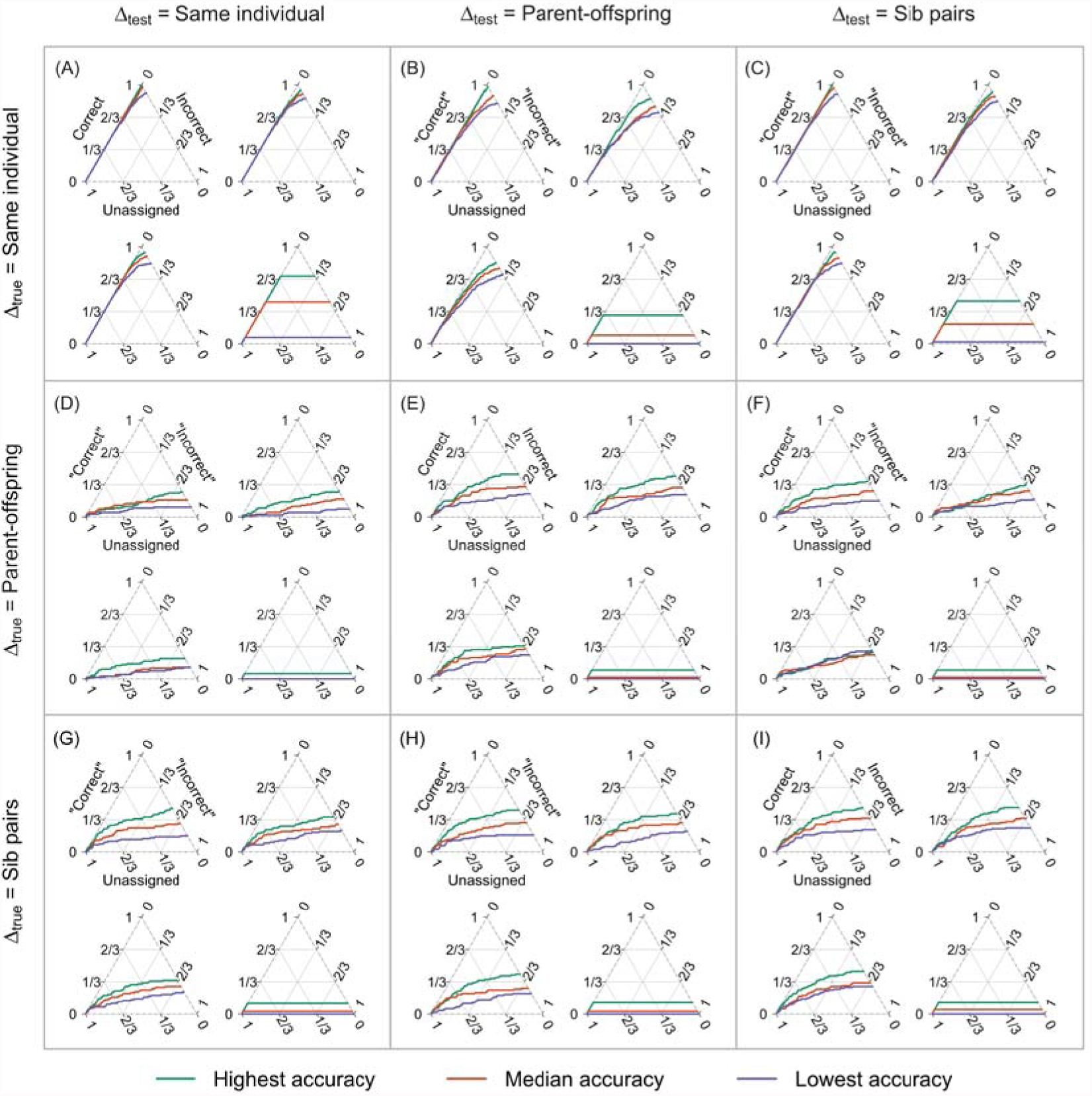
Proportions of profiles correctly assigned, incorrectly assigned, and unassigned with a varying match-score threshold. For each group of four plots, four match-assignment scenarios are shown: one-to-one matching (top left), one-to-many matching querying a SNP profile and selecting an STR profile with the highest match score (top right), one-to-one matching querying an STR profile and selecting a SNP profile with the highest match score (bottom left), and needle-in-haystack matching, counting the proportion of true matches with match scores exceeding the match scores of all non-matching pairs (bottom right). In each triangle, when the threshold is large, all profiles are unassigned (lower left vertex). Lowering the threshold leads to assignment of all profiles, tracing a curve to the right edge of the triangle. In needle-in-haystack matching, all putative matches have greater match scores than all putative non-matches; thus, once the match-score threshold falls below the largest match score among true non-matches, the number of correct matches remains constant, and the number of incorrect assignments increases while the number of unassigned profiles decreases. For each match-assignment scenario, we plot results from partitions with the minimum, median, and maximum record-matching accuracy across 100 partitions of the sample into training and test sets. (A) **Δ**_true_, same individual; **Δ**_test_, same individual. (B) **Δ**_true_, same individual; **Δ**_test_, parent–offspring. (C) **Δ**_true_, same individual; **Δ**_test_, sib pairs. (D) **Δ**_true_, parent–offspring; **Δ**_test_, same individual. (E) **Δ**_true_, parent–offspring; **Δ**_test_, parent–offspring. (F) **Δ**_true_, parent–offspring; **Δ**_test_, sib pairs. (G) **Δ**_true_, sib pairs; **Δ**_test_, same individual. (H) **Δ**_true_, sib pairs; **Δ**_test_, parent–offspring. (I) **Δ**_true_, sib pairs; **Δ**_test_, sib pairs. In panels (A), (E), and (I), **Δ**_true_ = **Δ**_test_. Correct pairs are matched with their true relationship, and incorrect pairs are unrelated but erroneously matched as related. In panels (B), (C), (D), (F), (G), and (H), **Δ**_true_ ≠ **Δ**_test_. “Correct” pairs are true relatives, but the hypothesized relationship is incorrect; “incorrect” pairs are non-relatives inferred to have the relationship in the test hypothesis.

When **Δ**_true_ = **Δ**_test_ = (0, 0, 0, 0, 0, 0, 1, 0, 0), in the median-accuracy partition in one-to-one matching, as the threshold is decreased, 164 of 218 (75.2%) profiles are correctly matched before an incorrect match is made (top left plot in Figure 3A). The corresponding values are 113 (51.8%) for the minimum-accuracy partition and 218 (100%) for the maximum-accuracy partition. With a decreasing threshold, the minimum-, median-, and maximum-accuracy partitions accurately match 106 (48.6%), 126 (57.8%), and 149 (68.3%) of 218 query SNP profiles (top right plot in Figure 3A), and 112 (51.4%), 154 (70.6%), and 184 (84.4%) of 218 query STR profiles (bottom left plot in Figure 3A), respectively, before an incorrect assignment occurs. In needle-in-haystack matching, in which all matches are incorrect after the first incorrect match, the median partition has 43.1% (94 of 218) accuracy (bottom right plot in Figure 3A). The minimum and maximum accuracies are 6.4% (14 of 218) and 69.7% (152 of 218), respectively.

For the parent–offspring case **Δ**_true_ = **Δ**_test_ = (0, 0, 0, 0, 0, 0, 0, 1, 0), few pairs are correctly matched before the first incorrect match (Figure 3E). With a threshold that permits false positives, however, many relative pairs are matched correctly: in one-to-one matching, minimum, median, and maximum accuracy are 26 (23.9%), 34 (31.2%), and 48 (44.0%) of 109 profiles, respectively. In one-to-many matching with a SNP query, these values are 25 (22.9%), 33 (30.3%), and 46 (42.2%) of 109, and they are 24 (22.0%), 35 (32.1%), and 42 (38.5%) of 109 for query STR profiles. Needle-in-haystack matching ranges from a minimum of 0 correct matches to a maximum of 10 (9.2%), with a median of 2 (1.8%).

Similar values to the parent–offspring case are obtained for 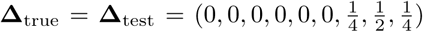, the sib-pair case (Figure 3I). Although even a stringent threshold produces false positives, many relationships are identified. In one-to-one matching, minimum, median, and maximum record-matching accuracy are 25 (22.9%), 38 (34.9%), and 50 (45.9%) of 109 profiles, respectively. Corresponding values are 27 (24.8%), 38 (34.9%), and 50 (45.9%) of 109 for one-to-many matching with query SNP profiles, and 28 (25.7%), 39 (35.8%), and 49 (45.0%) of 109 for query STR profiles. For needle-in-haystack matching, the minimum, median, and maximum are 0, 5 (4.6%), and 13 (11.9%) correct matches of 109, respectively.

As was observed in one-to-one matching, comparing corresponding panels within the rows of Figure 3, in one-to-many matching with a query SNP profile, one-to-many matching with a query STR profile, and needle-in-haystack matching, the record-matching accuracy with a misspecified test hypothesis is smaller than that seen with the correctly specified hypothesis (Figure 3B,C,D,F,G,H). Corresponding minimum, median, and maximum accuracies are similar under the misspecified hypothesis, as are the trajectories obtained as the match-score threshold decreases.

### Additional STRs

We evaluated the dependence of the record-matching accuracy on the number of STR loci by repeating our analyses with random sets of non-CODIS STRs. For each of the three choices of **Δ**_true_, considering the median-accuracy partition depicted in Figure 2 (panels A, E, I), we examined the record-matching accuracy for 100 randomly chosen sets of *L* loci, with *L* = 5, 10, 15*,…,* 100 (Methods, “Additional loci”).

For each pair (**Δ**_true_, **Δ**_test_) and each of the four match-assignment algorithms, Figure 4 depicts the median record-matching accuracy across the 100 locus sets. A comparison of panels within rows of the figure finds that record-matching accuracy is greater for one-to-one matching than for the two one-to-many matching scenarios, which in turn have higher accuracy than the needle-in-haystack scenario. In all panels, accuracy increases with the number of loci, nearing 100% when examining one-to-one and one-to-many matching with 100 loci, and exceeding 80% for needle-in-haystack matching with **Δ**_true_ = **Δ**_test_.

**Figure 4:**
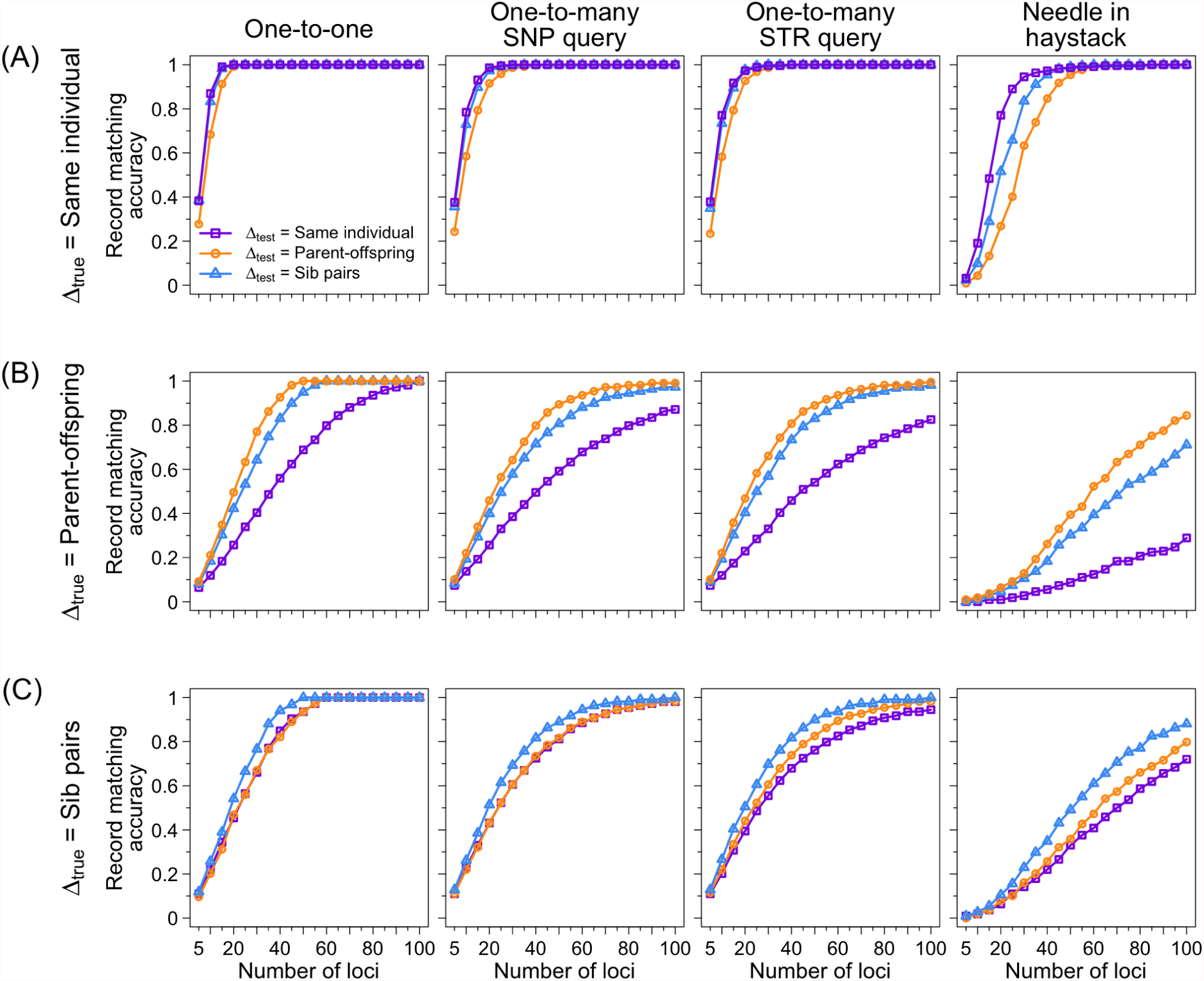
Record-matching accuracy as a function of the number of STRs. For each number of loci at intervals of 5 loci, 100 random locus sets are analyzed for the data partition in Figure 2. (A) **Δ**_true_, same individual. (B) **Δ**_true_, parent–offspring. (C) **Δ**_true_, sib pairs. In each of the 12 plots representing a choice of **Δ**_true_ and a match-assignment algorithm, three lines are shown, representing the median record-matching accuracy across 100 loci for each of three choices of **Δ**_test_.

Comparing panels within columns, accuracy is greater for SNP and STR profiles in which **Δ**_true_ represents exact matches (Figure 4A) than in cases with parent–offspring (Figure 4B) and sib-pair (Figure 4C) relationships. The correctly specified hypothesis produces greater accuracy than the two misspecified hypotheses, with the highest accuracy occurring for the same-individual hypothesis for **Δ**_test_ in the panels of Figure 4A, for the parent–offspring hypothesis in Figure 4B, and for the sib-pair hypothesis in Figure 4C.

## Discussion

We have found that not only can STR and SNP records be identified as belonging to the same individual, in many cases, STR and SNP profiles can be identified as belonging to close relatives—even though the profiles have no markers shared in common (Table 1). In one-to-one and one-to-many matching, record-matching accuracies were 30-32% for identification of parent–offspring pairs and 35-36% for identification of sib pairs, increasing toward 100% as the number of markers in the STR profile was increased to 100 (Figure 4).

The record-matching accuracies for parent–offspring pairs and sib pairs—relationships with the same overall kinship coefficient—were lower than the accuracies observed for STR and SNP profiles originating from the same individual. Record-matching of profiles from relatives is weakened because STR alleles in one profile of a pair and the neighboring SNP haplotypes in the other profile need not have been co-inherited from the same ancestor. Accuracies were slightly higher for sib pairs than for parent–offspring pairs.

Interestingly, when the relatedness hypothesis tested was misspecified, the reduction in accuracy was relatively small. Parent–offspring pairs were identified by record matching with a sib-pair hypothesis for the relationship with accuracy 27-28% for one-to-one and one-to-many matching, and sib pairs were identified with a parent–offspring hypothesis with 30% accuracy (Table 1). Pairs of relatives were also revealed by record matching when searching for exact matches, though with slightly lower accuracies of 17-18% for parent–offspring pairs and 28-29% for sib pairs. These results suggest that in practical settings in which the true relationship between profiles of interest to match is unknown, relative pairs will often be identified even when testing an incorrect relationship hypothesis.

This study contributes to a growing body of work on inference of genetic relationships in scenarios more challenging than when relatives are typed for the same markers [22-26]. In the setting of ancient DNA, Vohr et al. [22] focused on the scenario in which DNA sequence is generated for different DNA samples possibly representing the same or related individuals—from a burial site, for example—but sequence is sufficiently sparse that reads do not necessarily overlap between samples. In a computation focused on detecting samples representing the same individual, Vohr et al. [22] could distinguish simulated parent–offspring pairs and sib pairs from unrelated pairs; this computation amounts to demonstrating that under a same-individual hypothesis for **Δ**_test_, pairs with a parent–offspring or sib relationship for **Δ**_true_ were uncovered. A second study formally estimating relatedness from sparse sequence data while making use of LD between sites with data available in different sampled individuals was able to identify second- and third-degree relative pairs [24].

The potential to perform familial searching of forensic STR profiles in SNP databases generates both opportunities and privacy risks. Because accuracy was 30-36% for identification of first-degree relatives rather than above 90% as was seen for paired profiles from the same individuals, identification of relatives by record matching with our one-to-one and one-to-many algorithms will be possible in fewer cases. However, if a match to a particular query STR profile is of interest, an algorithm can be envisioned in which multiple top hits in SNP databases are further explored—by additional genotyping of contributors for whom DNA is available, or by genealogical tracing of relatives of the contributors to uncover exact matches. A relaxed accuracy measure in such a setting could therefore tabulate true matches that have high match scores (but not necessarily the highest value) or that differ in their scores from the highest value by less than a specified constant. Accuracy can also increase substantially even with small increases in the number of STR loci considered (Figure 4).

Record linkage of pairs of relatives between STR and SNP databases has a significant impact on genetic privacy. In addition to magnifying the exposure of the relatives of SNP-profile contributors to forensic identification, it also increases the phenotypic reach of STR profiles. CODIS genotypes of one individual could potentially be associated with genomic SNP genotypes of a relative, which could, in turn, reveal phenotypes of that relative [27]. Thus, not only could a CODIS genotype profile reveal phenotypic information about an individual [13], it could also reveal phenotypes of relatives. With access to SNP databases, the information contained in a CODIS profile would extend far beyond its value for identification of its contributor to also include genome-wide genetic data and phenotypic information about relatives of that contributor.

The possibility of performing familial searching of forensic profiles in SNP databases, while raising new concerns, also alters an existing concern, namely the unequal representation of populations in forensic databases. In profile queries to search for a relative already in a forensic database, populations overrepresented in the databases owing to overrepresentation in criminal justice systems are likely to produce more identifications of relatives, potentially contributing to further overrepresentation [5, 8, 9]. Record-matching queries to biomedical, genealogical, or personal-genomic databases, however, will instead produce more identifications in different sets of populations emphasized in genome-wide association studies and personal genomics data [9, 28, 29].

We note that we have not taken into account information on population of origin of the individuals; although the effect of population of origin in record-matching in the same-individual scheme was limited [13], relatedness profiling has been seen to produce varying accuracy by population [7, 8] and might do so in record-matching as well. The performance of the method will likely decrease with larger test sets numbering in the thousands or millions. In larger samples, the method will benefit from increased accuracy in inferring the LD pattern in the training set. However, increased sizes of test sets will reduce accuracy by generating larger numbers of possible matches. Finally, although we have focused on parent–offspring and sib-pair relationships, our framework to arbitrary relationship hypotheses more generally. For more distant relationships, however, accuracy will be lower in the same manner that it is reduced when comparing parent–offspring and sib-pair schemes to the case of matching profiles from the same individual. Nevertheless, this study contributes to growing understanding of the extent of the information contained in individual genotype profiles when those genotypes are analyzed together with databases of genotypes of other individuals, finding that that information can be considerable, both about the individuals typed and about their relatives.

## Methods

### Data

The data consisted of diploid genotypes of 872 individuals typed for 642,563 autosomal SNP loci, 13 autosomal STR loci from the CODIS panel, and 431 additional autosomal tetranucleotide STR loci. We used the same datasets as Edge et al. [13], containing no pairs of close relatives.

### BEAGLE settings

Inference of haplotype phase and imputation of STR genotypes were performed using `BEAGLE` v4.1 [19, 20]. For each STR marker, we considered in `BEAGLE` 1-Mb SNP windows extending 500 kb in each direction from the STR midpoint.

For phasing, which we performed in two different steps—one prior to pedigree generation, and another for phasing training sets—we used the default number of 10 iterations in `BEAGLE`, and we used default `BEAGLE` phasing parameters: maxlr=5,000, lowmem=false, window=50,000, overlap=3,000, impute=true, cluster=0.005, ne=1 million, err=0.0001, seed=-99,999, and modelscale=0.8.

We also used `BEAGLE` for imputation in test sets using phased training sets as reference data, employing a linkage map based on GRCh36 coordinates and the same parameters as in phasing, except gprobs=true and maxlr=1 million.

### Pedigree generation

Prior to generating pedigrees, we first phased the entire dataset (“`BEAGLE` settings”) to obtain individual haplotypes for use in pedigree generation. Next, we chose pairs of individuals without replacement to obtain 436 parental pairs from the 872 sampled individuals. For each parental pair, we simulated two offspring. We assumed that our 1-Mb window size was small enough that no recombination occurred within windows, so that haplotypes were transmitted intact in nuclear pedigrees. STR loci were treated as independent, so that assortment was independent across STRs.

Once pedigrees were generated, we dephased haplotypes, randomizing allele orders within individuals to hide phase information. We generated 10 sets of random pedigrees, each containing a distinct pairing of the 872 individuals into parental pairs. We used the same 10 sets in the parent–offspring and sib-pair schemes.

### Training and test sets

Following Edge et al. [13], we partitioned our dataset into a training set with 75% of the individuals and a test set of size 25%. For our “same individuals” computations, 654 of 872 individuals were assigned to the training set, and the other 218 to the test set. For these computations, we generated 100 random training-test partitions. We then imputed STR genotypes in the test set with the training set as a reference (“`BEAGLE` settings”), constructing a 218 × 218 match-score matrix to match imputed STR genotypes in the test set to actual test-set STR genotypes.

For the parent–offspring scheme, without loss of generality, we placed parent genotypes in the SNP dataset and offspring genotypes in the STR dataset. For each of the 10 sets of random pedigrees, we generated 10 random partitions into training and test sets, for a total of 100 training-test partitions. Each partition contained a training set of 75% of the pedigrees (327 pedigrees) and a test set with the other 25% (109 pedigrees). For each partition, we phased the training set (“`BEAGLE` settings”), including only the 654 parental individuals in the computation, to produce a phased reference panel. To produce a test set, from each test-set pedigree, we chose one parent and one offspring, creating 109 parent–offspring pairs. We then imputed STR genotypes of parents in the test set with the training set as a reference (“`BEAGLE` settings”) and constructed a 109 × 109 match-score matrix to match the imputed STR genotypes of parents in the test set to STR genotypes of offspring in the test set.

For the sib-pair scheme, as in the parent–offspring scheme, we generated 10 random partitions of each of the 10 sets of random pedigrees to produce 100 training-test partitions. For each partition, we assigned 75% of the pedigrees to the training set and 25% to the test set. We then took the parents from each training-set pedigree to form a training set that we then phased (“`BEAGLE` settings”). For a test set, a set of 109 siblings, each randomly selected from a pedigree, was used for the SNP dataset, and the remaining 109 siblings acted as an STR dataset. We imputed STR genotypes of the sibs in the test set with the training set as a reference (“`BEAGLE` settings”) and constructed a 109 × 109 match-score matrix to match the imputed STR genotypes in the half of the sibs treated as having SNP data in the test set to the STR genotypes of the other half of the sibs in the test set.

In selecting median-accuracy partitions, we chose the lesser of two possible median values among 100 partitions.

### Additional STRs

We scanned numbers of loci from 5 to 100 with an increment of 5. From 431 non-CODIS STR loci, we initially generated 100 sets of 100 random loci. For each set of loci, we recursively selected subsequent sets of loci with fewer loci at random so that the newly selected set of loci was nested in the previous set.

For each of the three true relationships (same individual, parent–offspring, sib pairs), we selected a partition and a pedigree set corresponding to the median one-to-one record-matching accuracy in the case of **Δ**_true_ = **Δ**_test_ (Figure 2A,E,I). We then ran our record-matching computations for each of the 100 sets of non-CODIS STR loci, considering each of the three test hypotheses (same individual, parent–offspring, sib pairs) and each of the four match-assignment scenarios (one-to-one, one-to-many with SNP query, one-to-many with STR query, needle-in-haystack).

### Match score calculation

To calculate the match score of Eq. 1 under an arbitrary relatedness hypothesis **Δ**, we must calculate the probability of STR profile *R_A_* given SNP profile *S_B_* and the relatedness hypothesis *M* = **Δ** for individuals *A* and *B*, 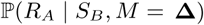, and the unconditional probability of *R_A_*, 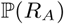.

Assuming independence of STR loci, the probabilities of STR probabilities are obtained as products across loci,

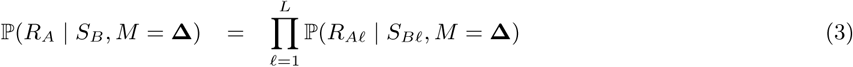

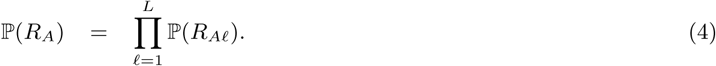

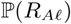 is calculated as the Hardy-Weinberg frequency of genotypes at locus *ℓ*.

To evaluate 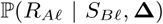 in Eq. 3, the probability for an individual *A* to have STR genotype *R_Aℓ_* at locus *ℓ*, conditional on a relative *B* having SNP genotype *S_Bℓ_* and relationship **Δ** = (Δ_1_, Δ_2_*,…,*Δ_9_), we sum over all values of *R_Bℓ_* in 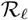, the set of all possible unordered diploid genotypes at STR locus *ℓ*:

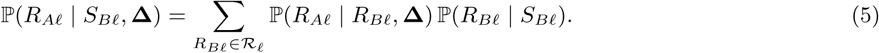

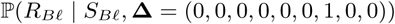, or 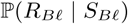 for short, is the probability for an individual *B* with SNP genotype *S_Bℓ_* at locus *ℓ* to have STR genotype *R_Bℓ_* at locus *ℓ*. We used `BEAGLE` to estimate the imputation probability for STR genotype *R_Bℓ_* given surrounding SNP profile *S_Bℓ_* in the same individual (see “`BEAGLE` settings”).

To obtain 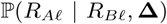 for arbitrary **Δ**, we denote the unordered STR genotypes of individuals *A* and *B* at locus *ℓ* by *R_Aℓ_* = *a_m_a_n_* and *R_Bℓ_* = *a_r_a_t_*, respectively. Alleles *a_m_*, *a_n_*, *a_r_*, and *a_t_* have allele frequencies *p_m_*, *p_n_*, *p_r_*, and *p_t_*, respectively, and they are not necessarily distinct. We decompose 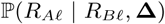 by conditioning on all possible condensed identity states *C_k_* describing the pair of individuals *A* and *B* (Table 1):

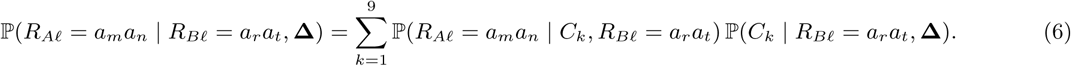

To evaluate the summands, we consider two separate cases: (1) individual *B* is heterozygous, and (2) individual *B* is homozygous. In each case, we assume Hardy-Weinberg genotype frequencies at locus *ℓ*.

### Individual *B* is heterozygous: *a_r_* ≠ *a_t_*

Because *B* is heterozygous, condensed identity states *C*_1_, *C*_2_, *C*_3_, *C*_4_, all of which assume identity by descent for the pair of alleles in individual *B*, are not possible. Thus, using the joint probability distribution of *C_k_* and the genotype of the individual *B* (Table 1), a state *C_k_* for *k* ≥ 5 has probability:

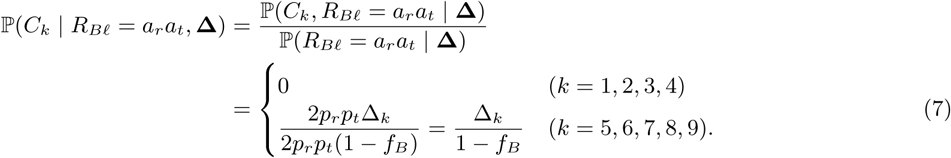

Here, because 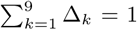, we simplify notation by using the inbreeding coefficient of individual *B*, or *f_B_* = Δ_1_ + Δ_2_ + Δ_3_ + Δ_4_.

To evaluate Eq. 6, it remains to compute 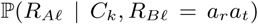 for each condensed identity state *C_k_*, *k* = 5, 6, 7, 8, 9. For the remainder of the case of *B* heterozygous, we treat *m*, *n*, *r*, and *t* as distinct.

***C*_5_**: Because the two alleles of *A* are identical by descent (IBD), *A* is homozygous. These alleles are IBD with one of the alleles of *B*, so *a_m_a_n_* = *a_r_a_r_* or *a_m_a_n_* = *a_t_a_t_*. Because the two alleles of *B* have equal probability of being IBD with the alleles of *A*, *R_Aℓ_* = *a_r_a_r_* and *R_Aℓ_* = *a_t_a_t_* are equally probable. Thus,

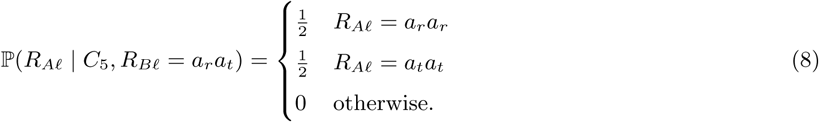

***C*_6_**: Because the two alleles of *A* are IBD, *A* is homozygous. However, the alleles of *A* are not IBD with any alleles of *B*, so *aman* can be any homozygous genotype in the population:

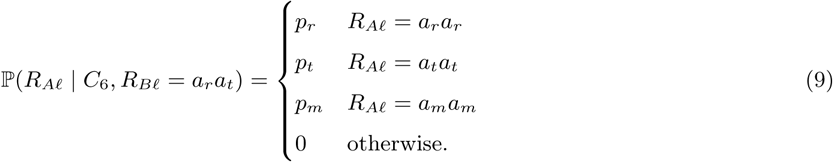

***C*_7_**: One allele of *A* is IBD with one allele from *B*, and the other allele from *A* is IBD with the other allele from *B*. Thus, *A* and *B* have the same unordered genotype: *a_m_a_n_* = *a_r_a_t_*:

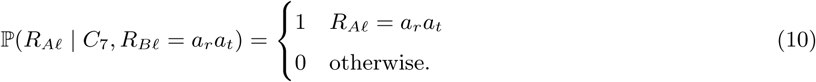

***C*_8_**: The only identity-by-descent relationship is between one allele of *A* and one allele of *B*. Thus, *A* can have genotype either *a_r_a_v_* or *a_t_a_v_*, where *a_v_* can be any allele in the population. Because *a_r_* and *a_t_* have the same probability of being the allele of *B* that is IBD with an allele in *A*, we have:

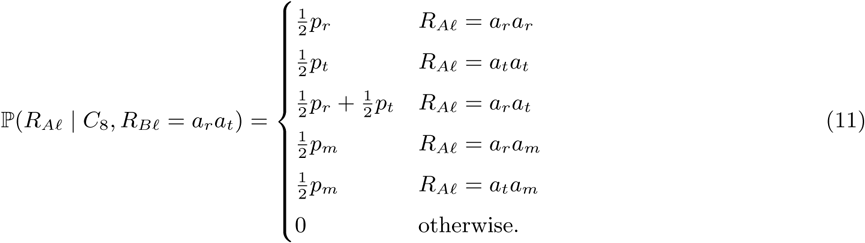

***C*_9_**: No alleles in *A* are IBD with an allele in *B*. Hence, genotype probabilities in *A* follow Hardy-Weinberg frequencies:

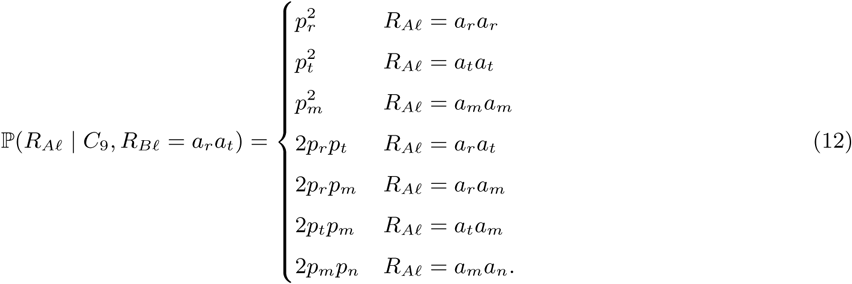

Combining Eqs. 7-12, for heterozygous *R_Bℓ_*, with *r*, *t*, *m*, and *n* all distinct and *R_Bℓ_* = *a_r_a_t_*, we have:

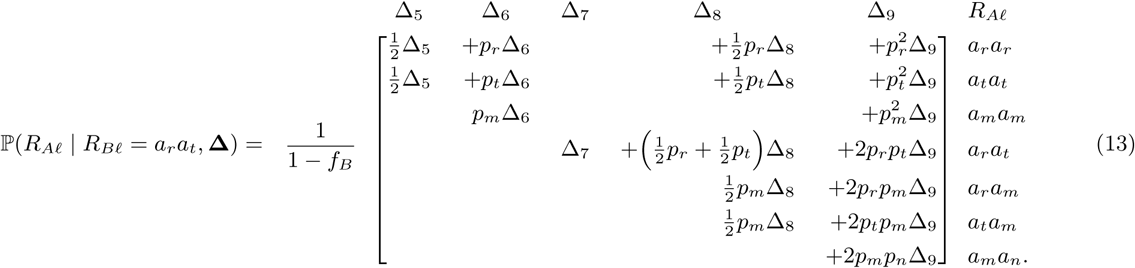

In this matrix notation, each row represents a probability for a particular choice of *R_Aℓ_*, summing across cases ***C*_5_**, ***C*_6_**, ***C*_7_**, ***C*_8_**, and ***C*_9_**. The distinct alleles *a_m_* and *a_n_* refer to alleles different from *a_r_* and *a_t_*. We use notation *T_rt_*(*x, y*) to represent the quantity associated with *R_Aℓ_* = *a_x_a_y_* in Eq. 13.

### Individual *B* is homozygous: *a_r_* = *a_t_*

When *B* is homozygous with genotype *a_r_a_r_*, all nine condensed identity states are possible. Using Table 1, the condensed identity state *C_k_* has probability:

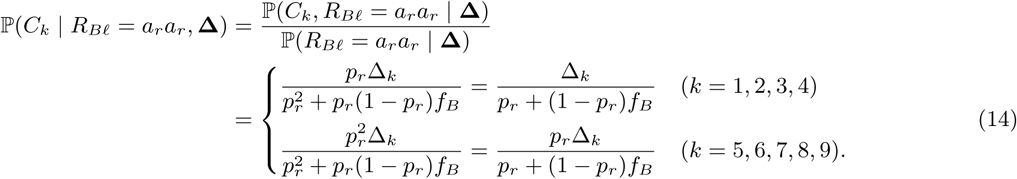

***C*_1_**: All four alleles from *A* and *B* are IBD: *a_m_* = *a_n_* = *a_r_* = *a_r_*. The genotype of *A* is fully specified by the genotype of *B*:

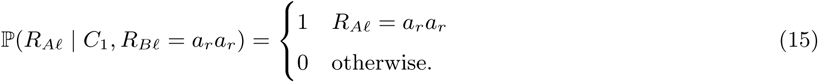

***C*_2_**: Alleles are IBD within individuals, but not between individuals: *a_m_* = *a_n_* and *a_r_* = *a_t_*. Hence, *A* must be homozygous, but not necessarily for the same allele as *B*. The probability for *A* to have homozygous genotype *a_m_a_m_* is the frequency of allele *a_m_*:

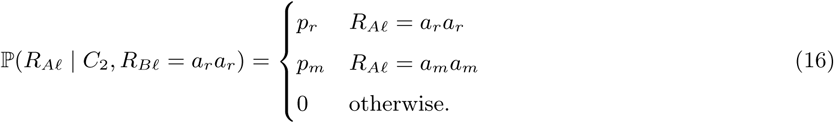

***C*_3_**: One of the alleles of *A* is IBD with both alleles of *B*. Because *B* is homozygous with genotype *a_r_a_r_*, the genotype of *A* is *a_r_a_v_*, where *a_v_* is any possible allele.

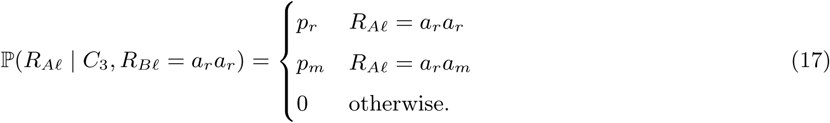

***C*_4_**: Both alleles in *B* are IBD but no identity by descent occurs for *A*. Thus, *A* can have any genotype in the population:

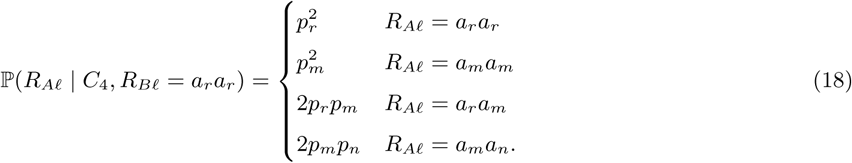

***C*_5_**: Both alleles in *A* are IBD, so *A* is homozygous. Because both alleles in *A* are IBD with one of the alleles in *B* and because *B* is homozygous with *a_r_a_r_*, the only possible genotype of *A* is *a_r_a_r_*:

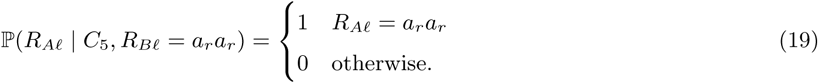

***C*_6_**: Both alleles in *A* are IBD and *A* is homozygous, but no identity by descent occurs with *B*. Thus, *A* can have any homozygous genotype:

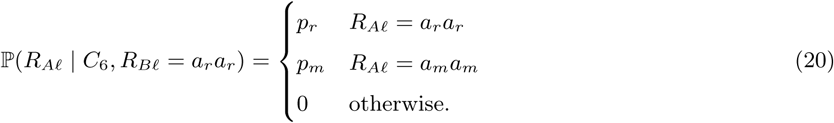

***C*_7_**: One allele in *A* is IBD with one allele in *B*, and the other allele in *A* is IBD with the other allele in *B*. *A* therefore has the same genotype as *B*:

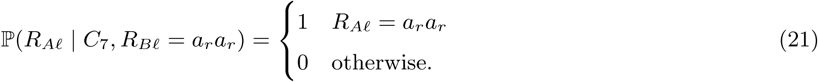

***C*_8_**: One allele of *A* is IBD with one allele of *B*, and remaining alleles of *A* and *B* have no identity by descent. Because *B* is homozygous with genotype *a_r_a_r_*, *A* has genotype *a_r_a_v_*, where *a_v_* is any possible allele:

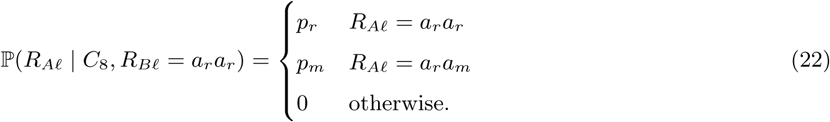

***C*_9_**: None of the alleles are IBD, and *A* can have any possible genotype:

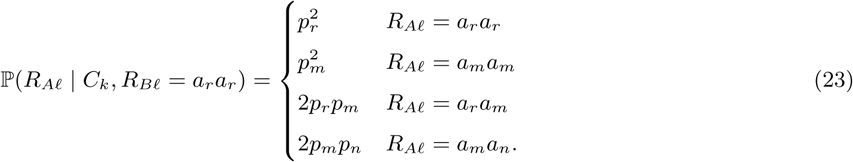

Combining all nine cases, from Eq. 14-23, with *a_m_*, *a_n_*, and *a_r_* all distinct and *R_Bℓ_* = *a_r_a_r_*, we obtain:

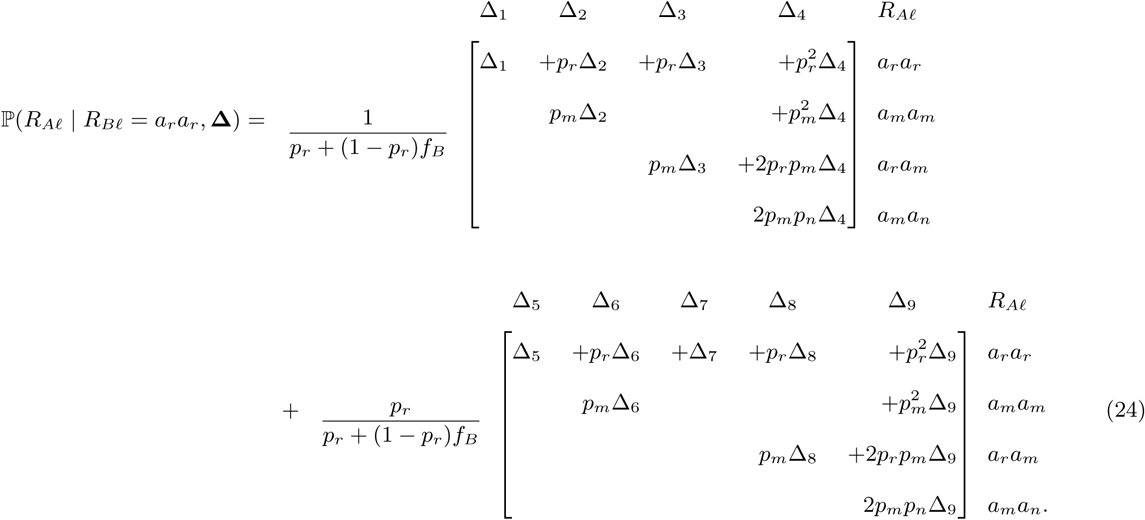

Here, the alleles *a_m_* and *a_n_* are distinct and indicate any alleles different from *a_r_*. We use notation *H_rr_* (*x, y*) to represent the quantity associated with *R_Aℓ_* = *a_x_a_y_* in Eq. 24.

### Completing the calculation

We can now expand Eq. 3 for arbitrary relationships *M* = Δ. Let *N_ℓ_* denote the number of distinct alleles possible at STR locus *ℓ*, and index these alleles by {*a*_1_*, a*_2_*,…, a_Nℓ_*}. From Eqs. 5, 13, and 24, recalling that 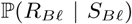 is obtained from `BEAGLE`, we have

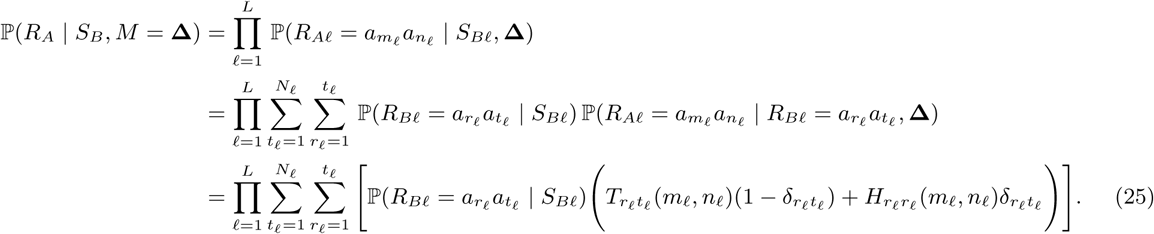

In the last step, we use the Kronecker delta to combine the heterozygous case of Eq. 13 and the homozygous case of Eq. 24 into a single equation.

## Acknowledgments

We acknowledge support from National Institutes of Health grant R01 HG005855 and National Institute of Justice grant 2014-DN-BX-K015.

